# Four new Duchenne muscular dystrophy mouse models with clinically relevant exon deletions in the human *DMD* gene

**DOI:** 10.1101/2025.11.21.689489

**Authors:** Maaike van Putten, Margot Linssen, Christa Tanganyika-de Winter, Conny M. Brouwers, Jill W.C. Claassens, Nisha Verwey, Max Walsh, Tiberiu Loredan Stan, Annemieke Aartsma-Rus, Peter Hohenstein

**Author notes:** **Corresponding authors:** Annemieke Aartsma-Rus –, Peter Hohenstein –.

## Abstract

Mutation specific therapeutic approaches, like exon skipping or gene-editing, hold promise for the treatment of Duchenne muscular dystrophy (DMD). Translatability of preclinical studies investigating these approaches could greatly be improved through the use of humanized mouse models, as these allow preclinical testing of human specific sequences.

We developed four novel humanized DMD mouse models with either a deletion of exon 44, 45, 51 or 53 in the human *DMD* gene, in a mouse dystrophin negative background (*mdx* mouse; exon 23 nonsense mutation). Our optimized prescreening pipeline allowed us to do so very efficiently with the CRISPR-Cas9 technology. We confirmed either complete lack of dystrophin, or expression of trace levels, which led to development of muscle pathology consisting of muscle fiber de-, and regeneration, inflammation and fibrosis in young adult mice. Intramuscular treatment with vivo-morpholinos targeting a flanking exon induced exon skipping in the DMD strains, which restored the disrupted open reading frame and subsequently dystrophin expression. This validates these models as valuable tools for preclinical studies investigating human sequence specific therapeutic approaches for DMD.

**Summary statement:** Humanized Duchenne muscular dystrophy mouse models were created with deletions of exon 44, 45, 51 or 53 in the human *DMD* gene. These dystrophic models allow preclinical testing of human-specific dystrophin restoring approaches.

## Introduction

Duchenne muscular dystrophy (DMD) is an X-linked, progressive, muscle-wasting disease that is caused by pathogenic variants in the dystrophin protein encoding *DMD* gene (Muntoni, Torelli, and Ferlini 2003). Normally, the dystrophin protein connects the cytoskeleton of muscle fibers to its connective tissue surrounding, by connecting F-actin and ß-dystroglycan with its N- and C-terminal domains, respectively. Loss of dystrophin protein makes muscle fibers vulnerable for contraction induced damage, leading to chronic inflammation, failure to regenerate and eventually replacement of muscle tissue by fibrotic and adipose tissues. Patients irreversibly lose muscle function, and even with multidisciplinary care they generally lose ambulation in their early teens, need assisted ventilation around the age of 20 and die in the 2^nd^-4^th^ decade of life due to respiratory or heart failure (Duan et al. 2021).

Whereas DMD patients carry pathogenic variants that prevent production of functional dystrophins, variants that do not disrupt the open reading frame and are located in the center of the gene allow the production of partially functional dystrophin proteins. These variants cause the allelic disease called Becker muscular dystrophy (BMD). This disease generally has a later onset and a slower disease trajectory compared to DMD. For both DMD and BMD, most patients carry large deletions involving one or more exons, which cluster between exon 42 and 55 (Bladen et al. 2015).

Therapeutic approaches aiming to restore dystrophin for DMD are based on the discrepancy between DMD and BMD mutations. Exon skipping and gene editing approaches aim to restore the open reading frame for *DMD* transcripts at the RNA and DNA level, using antisense oligonucleotides (ASOs) and CRISPR/Cas9, respectively. These approaches are variant specific, as based on the size and location of the mutation, different exons need to be skipped to restore the reading frame (Niks and Aartsma-Rus 2017; Roberts, Wood, and Davies 2023).

Animal models for DMD have been invaluable to study disease pathology. For therapeutic development, especially mouse models have been essential (van Putten et al. 2020; Zaynitdinova, Lavrov, and Smirnikhina 2021). The most commonly used mouse model for DMD is the *mdx* mouse (Bulfield et al. 1984) and standard operating procedures for its use in preclinical studies have been developed (Willmann, Dubach, and Chen 2011). This model carries a spontaneous nonsense mutation in exon 23 of the mouse *Dmd* gene (Sicinski et al. 1989). It originally arose on the C57BL/10ScSnJ background, but has since then been backcrossed to other backgrounds including C57BL/6J and DBA/2J (Fukada et al. 2010).

The *mdx* mouse has been used to provide proof-of-concept for the ability to restore dystrophin with exon skipping and gene editing approaches (Lu et al. 2003; Nelson et al. 2016). However, these studies focused on ASOs and guide RNAs targeting mouse dystrophin exon 23 and therefore were poorly translatable to the human situation. Firstly, different ASOs and guide RNAs were required as the mouse equivalents were not effective in humans due to species specific differences in sequences. Secondly, most DMD patients have pathogenetic variants in a different region of the gene and as such, the dystrophin protein produced by the *mdx* mouse after exon skipping or gene editing, differs from the ones that will be produced by the majority of patients. To allow testing of human specific ASOs or guide RNAs, a YAC transgenic mouse model carrying the complete 2.3 MB human *DMD* locus was generated (’t Hoen et al. 2008). This model carried the human *DMD* (hDMD) gene on mouse chromosome 5. The expression of human dystrophin protein was capable of rescuing the *mdx* phenotype. To facilitate preclinical studies with human specific ASOs or guide RNAs, models with pathogenic variants within the *hDMD* gene are required. We previously used TAL effector nucleases to delete human exon 52 from the *hDMD* gene as a first effort towards this goal (hDMDdel52/*mdx* strain) (Veltrop et al. 2018). In the course of validating this model it was found that the hDMD YAC model carries two copies of the YAC in a tail-to-tail orientation, where the hDMDdel52/*mdx* model carries a partial deletion of exon 52 on both of the hDMD copies (Yavas et al. 2020). This realization explained why generating the hDMDdel52/*mdx* model had failed using homologous recombination. Furthermore, the tail-to-tail orientation severely complicated the generation of additional deletion models with genome editors, as there was a risk of deleting the target exon (e.g. exon 45) but also the entire content in between (exon 45-79 from both copies of the YAC (Fig. 1)).

**Figure 1.**
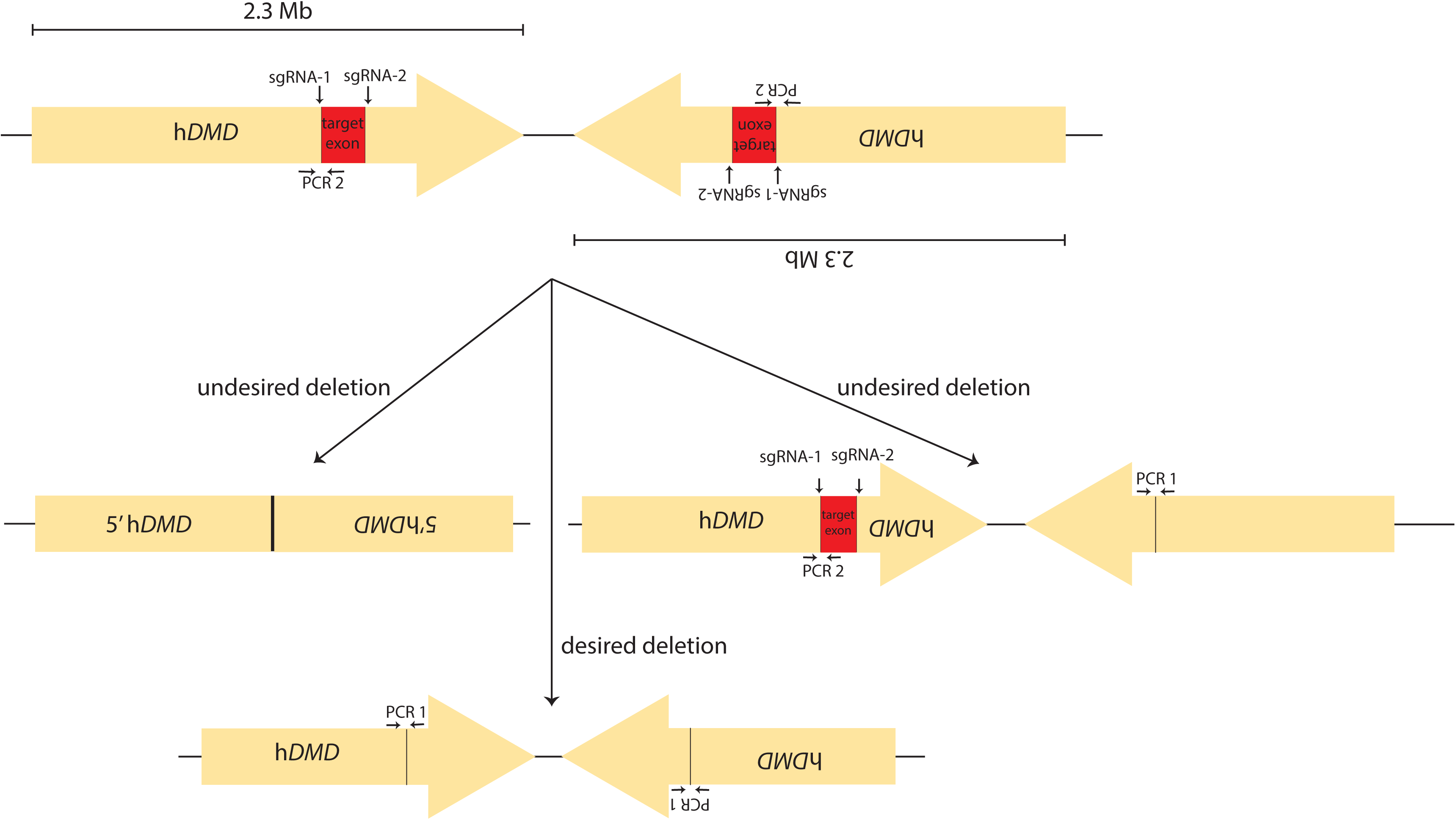
The hDMD YAC transgene locus and prescreen PCR design for generating new exon deletion mutants. Two PCR assays were sufficient to identify candidates for correct deletion of both copies of the target exon. PCR 1 gives a predictable fragment size after correct deletion by spanning the target exon. PCR 2 detects the retention of the target exon and the 5’ end of the *DMD* gene. The hDMD YAC transgenic mouse has two copies of the *DMD* YAC in a tail-to-tail configuration and scores negative for the predicted product size for PCR 1. Mice scoring positive for a PCR fragment of the predicted size for PCR 1 have lost at least one of the copies of the target exon and must have retained the 5’ half of both copies of the YAC. The mice were further tested for absence of PCR 2 which is specific for the undeleted target exons. Mice positive for PCR 1 and negative for PCR 2 were selected for further quality screening.

Here, we optimized the procedure to make additional clinically relevant models from the hDMD/*mdx* mouse and used this to generate four new DMD mouse models carrying a deletion of either exon 44, 45, 51, or 53 in the hDMD gene in the C57BL/6J/*mdx* background. We further demonstrate dystrophin restoration after ASO-induced exon skipping for each of the models to validate them for preclinical testing of human-specific therapies.

## Materials and Methods

### Mouse lines

The following mouse lines were generated or used in this study. Tg(DMD)72Thoen/J (MGI:5578551); Dmd*mdx* (MGI:1856328); Tg(DMD*)del44Lumc (MGI:7408337); Tg(DMD*)del45Lumc (MGI:7408338); Tg(DMD*)del51Lumc (MGI:7408339); Tg(DMD*)del53Lumc (MGI:7408340). The animal care and experimental procedures were approved by the Animal Welfare Body Leiden and are in accordance with the Dutch Experiments on Animals Act and EU Directive 2010/63/EU.

### CRISPR/Cas9 editing

All CRISPR editing was done using RNP complexes made from Alt-R CRISPR recombinant Cas9 and synthetic crRNA and tracrRNA (IDT) assembled according to the suppliers’ protocol. All guide RNA sequences used are given in Supplementary Table 1.

### ES culture and blastocyst injection

hDMD/*mdx* ES cells have previously been described (Veltrop et al. 2013). To generate Tg(DMD*)del44Lumc, cells were grown on a monolayer of mitotically inactivated mouse embryonic fibroblasts (MEFs) in KnockOut DMEM supplemented with 2 mM L-Glutamine, 1 mM Sodium Pyruvate, 1x MEM Non-Essential Amino Acid, 0,1 mM 2-Mercaptoethanol, 1000 units/ml ESGRO Leukemia Inhibitory Factor and 10% Fetal Bovine Serum and cultured on 0,1% gelatine coated cell culture plates. For CRISPR-mediated editing of the ES cells we used two CRISPR RNAs annealed to Alt-R™ CRISPR-Cas9 tracrRNA. The RNP complex was made using 50% Alt-R™ S.p. dCas9 Protein V3 and 50% Alt-R™ S.p. Cas9 Nuclease V3 according to the IDT protocol. The RNP complex was electroporated into the ES cells using the mouse embryonic stem cell nucleofector kit (Lonza) and the nucleofector II (Amaxa). Clones were picked into 96 well plates, expanded, and duplicated for freezing and DNA isolation for PCR prescreening; 30 sec 95°C, [20 sec 95°C, 20 sec 67°C, 40 sec 72°C for 35 cycli] 1 min 72°C, using dreamtaq polymerase. All prescreening primers are given in Supplementary Table 2. Primers in intron 43 and intron 44 resulted in a PCR product of ∼500 bp or smaller in case of a deletion of exon 44 (Fig. 2). Primers in exon 44 and intron 44 were used to check for the presence of exon 44, indicated by the absence of the ∼1600 bp fragment. Copy count by qPCR was done for human *DMD* exons 1, 43, 44, 45 and 79, and chromosomes where counted on candidate clones on metaphase spreads. Correctly targeted ES cells were injected into C57BL/6JrJ mouse blastocysts. The deletion, as determined in the targeted ES cells, was confirmed by ddPCR and Sanger Sequencing (Supplementary Tables 2 and 3). Copy count for all exons was done on the BIORAD QX200 Droplet Digital PCR System according to the protocol provided by the manufacturer. Probes were designed with IDT PrimerQuest™ (Supplementary Table 3). All assays were done in Biorad ddPCR™ Supermix for Probes (No dUTP) #1863023 with 750 µM of each primer, 250 µM of each probe and 50 ng DNA purified from ear or tail with Qiagen DNeasy Blood and Tissue Kit (Cat no. / ID. 69504). A single exon probe was amplified in parallel with a reference gene. After droplet generation on the QX200 Droplet Generator, a PCR was performed; 10 min 95°C, [30 sec 94°C, 1 min 60°C for 39 cycli] 10 min 98°C on a Biorad T100 Thermal Cycler, droplets where measured on the QX200 Droplet reader and results analyzed with QX200 Quantasoft Analysis Pro version 1.7.4.

**Figure 2.**
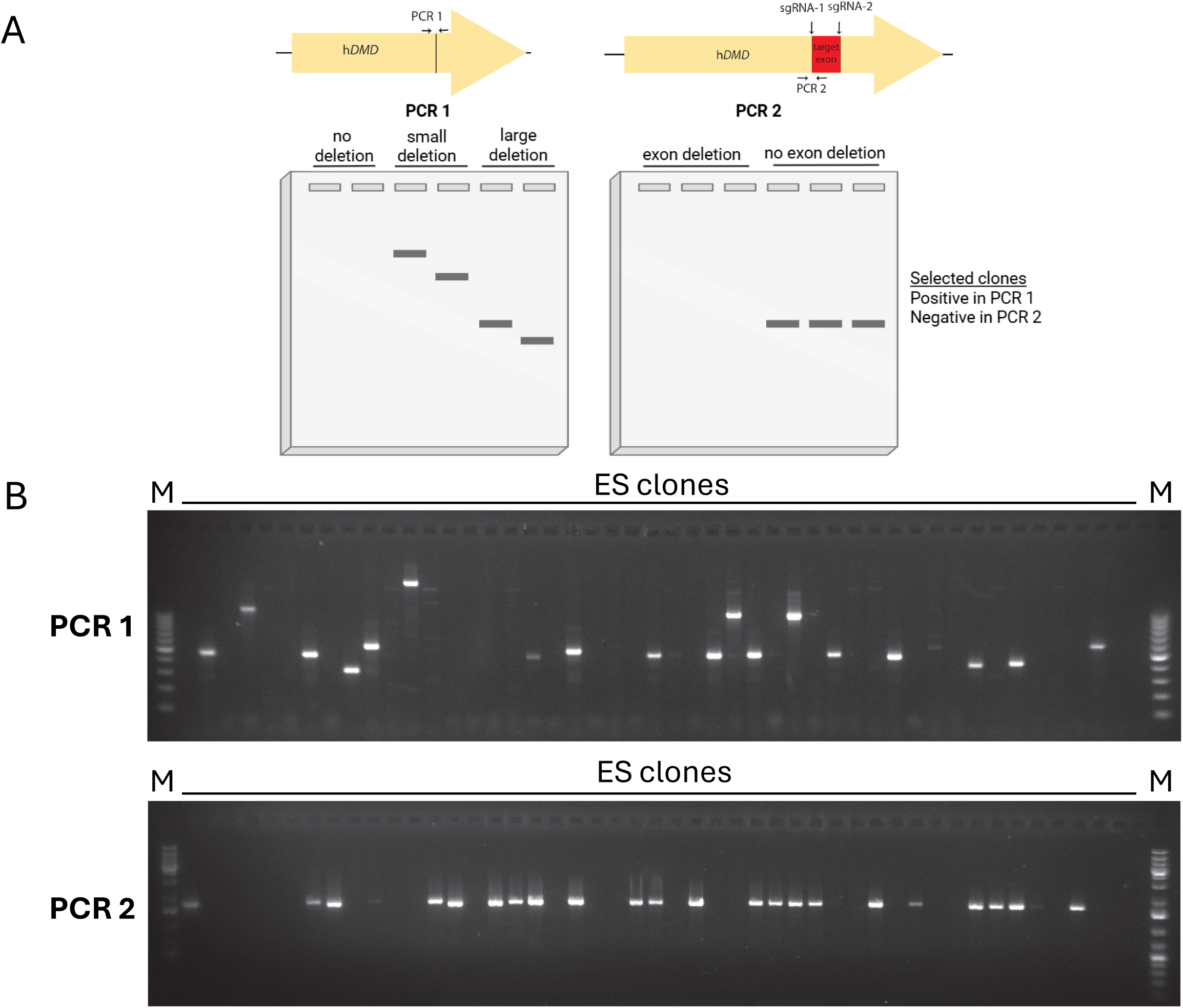
Prescreen PCR of hDMD/*mdx* ES cells targeted for deletion of exon 44. **A.** Illustration of expected PCR product sizes for PCR 1 performed after transfection, aimed to detect clones with a deletion of exon 44, in which the project size depends on the size of the deletion and PCR 2 performed on clones positive in PCR 1, aimed to check for the absence of the undeleted exon 44. **B.** PCR 1: Example of ES clones after transfection with CRISPR/Cas9 and single guide RNAs designed to delete hDMD exon 44. The PCR conditions were chosen for not being able to amplify the undeleted allele. PCR 2: Example of PCR assay on del44-positive ES clones to detect any remaining exon 44. Clones that scored positive for this assay would have lost only one copy of the exon and were discarded. Only clones positive for PCR 1 (panel A) and negative for PCR 2 were tested further. M; 100bp ladder.

Chimeric males were bred to C57BL/6Jico/Lumc to establish the Tg(DMD*)del44Lumc (hDMDdel44/*mdx*) mouse line. For Sanger sequencing (in general) the same primers were used as the ones used for PCR. Both a forward and a reverse read was performed on the PCR product to confirm results, which were analyzed with Snapgene software. Genotyping for the *mdx* exon 23 mutation was performed as previously described (Shin et al. 2011).

### Mouse zygote electroporation

The Tg(DMD*)del45Lumc (hDMDdel45/*mdx*), Tg(DMD*)del51Lumc (hDMDdel51/*mdx*) and Tg(DMD*)del53Lumc (hDMDdel53/*mdx*) lines were generated directly in embryos via electroporation at the zygote stage. For electroporation, a mix of components was prepared in Opti-MEM (Gibco; 31985062) including ctRNA 100ng/μl (TracrRNA and crRNA annealed in IDT duplex buffer), SpCas9V3 100ng/μl and ssODN 50ng/μl (IDT technologies). Freshly obtained fertilized C57BL/6JrJ zygotes were added to the mix and electroporation was performed using NEPA21 (NEPAGENE) in a CUY501P1-1.5 slide, with the following settings: Poring pulse (40 V, pulse duration 3ms, pulse interval 50 ms, 4 pulses) and transfer pulse (5 V, 50ms, 50ms, 5 pulses).

Deletion of both copies of exon 45 or 53 was confirmed in the mosaic offspring using ddPCR copy count and subsequent Sanger sequencing of the deletion-specific prescreen PCR fragments using the same primers as used for amplification. PCR conditions for exon 45; 10 min 95°C, [30 sec 95°C, 30 sec 60°C, 50 sec 72°C for 35 cycli] 5 min 72°C, using DreamTaq polymerase. PCR conditions for exon 53; 5 min 98°C, [10 sec 98°C, 10 sec 66°C, 20 sec 72°C for 30 cycli] 5 min 72°C, using Q5 polymerase (Supplementary Table 2 and 3). The exon 53 mutant was found to have two different deletions in the copies of the YAC. Here, we cloned the mixed PCR fragment in the pCR4-TOPO TA Vector (Thermo Fisher Scientific) according to the supplied protocol and sequenced 12 clones using the M13F and M13R primers in the vector backbone. For Tg(DMD*)del51Lumc we only obtained founder animals that had lost one copy of the hDMD YAC. Therefore, F1 offspring was subjected to a second round of electroporation with specific guide sequences located within the deleted region, resulting in a mouse line carrying both a deletion of both DMD copies, but a slightly different spanning region. Also for this strain, pups were checked for copy counts by ddPCR and Sanger sequencing as described above (Supplementary Table 2 and 3). Furthermore, they were genotyped with the same PCR conditions as those used for the exon 45 PCR. Genotyping for the *mdx* exon 23 mutation was performed as previously described (Shin et al. 2011).

### Histological assessments

Sections of 8µm thick were generated of the gastrocnemius of 8-weeks-old untreated hDMD/*mdx*, C57BL/6J, *mdx*, hDMDdel44/*mdx*, hDMDdel45/*mdx*, hDMDdel51/*mdx* and hDMDdel53/*mdx* males. Hematoxylin and Eosin staining was performed to visualize overall histopathology. After defrosting of the slides, they were fixated in ice cold acetone for 5 min and rinsed with water. Slides were incubated with Hematoxylin for 3 min, and the stain was developed with tap water for 5 min. Thereafter, slides were dipped 8-10 times in acid ethanol and submerged in eosin for 30 sec, followed by incubation in increasing concentrations of ethanol (80, 90, 100%) for 1 min each. After 5 min incubation in xylene, slides were mounted with Pertex and images were taken at a 20x magnification with the BZ-X710 all-in-One Fluorescence Microscope (Keyence, the Netherlands).

The same samples were also stained for dystrophin and laminin. Hereto, sections were defrosted and fixated with ice cold acetone for 5 min, washed for 1 min in PBS and blocked with 1x PBS Tween 20 (0.05%) with 5% horse serum. Slides were incubated with rabbit anti-dystrophin (1:250 Ab15277, Abcam) and rat anti-laminin α2 (1:50, SC-59854, Santa Cruz Biotechnology), in blocking buffer for 60 min. Thereafter, slides were incubated in blocking buffer with goat-anti-rabbit Alexa 594 (1:1000, A-11037, Thermo Fisher Scientific, the Netherlands) for dystrophin and goat-anti-rat Alexa 488 (1:1000, A-11006) for laminin. After washing, slides were mounted with ProLong™ Gold Antifade Mountant with DNA Stain with DAPI (Thermo Fisher Scientific). Pictures were made at a 20x magnification using the BZ-X710 all-in-One Fluorescence Microscope (Keyence, the Netherlands).

### Western blotting

Muscle sections were homogenized with protein isolation buffer (1.25 mM Tris-HCl pH 6.8, supplemented with 25% (w/v) SDS) for 20 sec at speed 7000 for 2-5 times with the MagNA Lyser and heated at 95°C for 10 min. The protein concentration was determined with the Bicinchoninic Acid Protein Assay Kit (Thermo Fisher Scientific) according to manufacturer’s instructions. Total protein (30µg) was added to Laemmli buffer (1.5M Tris pH 6.8, 10% v/v glycerol, 20% SDS) and ß-mercaptoethanol solution and heated at 95°C for 10 min. Samples were run on a 3-8% Tris-Acetate gel (Bio-Rad Laboratories) for 1 hour at 75V and for 1 hour at 150V. Blotting was performed with the Trans-Blot Turbo system with a midi nitrocellulose package at 2.5A for 10 min. Membranes were incubated with blocking buffer (5gr. non-fat dried milk powder (Sigma) in 100ml 1x TBS (Tris-Buffered Saline)) for 1 hour and washed three times 15 min with 1x TBST (TBS with Tween-20). Mouse and human dystrophin protein was detected using the Ab154168 primary antibody (1:2000, Abcam) in combination with the loading control α-actinin (1:1000, 66895-1, Proteintech). Human dystrophin was detected with the Mandys106 antibody (1:125, Sigma Aldrich) in combination with loading control α-actinin (1:1000, Ab72592, Abcam). Tris-Buffered Saline Incubations were performed in Takara Immuno booster 1 (T7111A, Takara, Göteborg, Sweden) at 4°C overnight. After 3x washing in TBST for 15 min, membranes were incubated with secondary antibodies IRDye 800 CW donkey-anti-rabbit (1:5000, Li-Cor) and IRDye 680RD donkey-anti-mouse (1:10,000, Li-Cor), targeting dystrophin and α-actinin, respectively. Membranes were washed in TBST and TBS (both 20 min) and analyzed with the OdysseyCLx imager and software (Li-Cor).

### Antisense oligonucleotide treatment

ASOs targeting exon 43, 44, 50 or 52 were designed (Supplementary Table 4) and ordered as vivo-morpholinos (ViM, Gene Tools, LLC). ViMs targeting exon 43 was used in the hDMDdel44/*mdx* model, exon 44 ViMs in the hDMDdel45/*mdx* model, exon 50 ViMs in the hDMDdel51/*mdx* model and exon 52 ViMs in the hDMDdel53/*mdx* model. Mice received intramuscular injections into the gastrocnemius (50µl) and triceps (40µl) with ViMs at a dose of 50µg (exon 50 and 52 ViMs), or 100µg (exon 43 and 44 ViMs) per muscle under isoflurane anesthesia on two consecutive days. Animals were sacrificed two weeks after the last injection and muscles were isolated, snap frozen in liquid nitrogen cooled isopentane and stored at - 70°C. Sections were generated with a cryotome and divided over two 1.4 mm Zirconium Beads Pre-Filled Tubes (OPS Diagnostics), to allow extraction of both RNA and protein from the same muscle sample.

### RNA isolation and exon skipping analysis

Muscles pieces were homogenized in TRIsure buffer (GC Biotech, the Netherlands) using the MagNA Lyser (Roche Diagnostics). Total RNA was isolated with chloroform in a 1:6 ratio on ice for 5 min. Samples were centrifuged (15,400 rcf for 15 min at 4°C) and the upper aqueous phase was transferred to a clean tube and an equal volume of isopropanol was added and precipitated for 30 min. After centrifugation (15,400 rcf for 10 min at 4°C), the pellet was washed with 70% ethanol and dissolved in RNase/DNase free water. For cDNA synthesis, 1000ng RNA was added to a mix of dNTPs (10mM) and random hexamer primers (40ng/µl) and incubated at 70°C for 5 min and chilled on ice. Thereafter, 5x reaction buffer (Promega), RNAsin (40U/µl, Promega), M-MLV reverse transcriptase (200U/µl, Promega) and MilliQ were added and left to incubate at 25°C for 10 min, 42°C for 60 min and 70°C for 10 min. Samples injected with saline, minus RT and water were included as negative controls. Per sample, 2.5µl cDNA was added to a PCR mixture consisting of 10x DreamTaq buffer, dNTPs (10mM), DreamTaq, MilliQ and ASO specific primers (ViMs targeting exon 43 or 44: forward primer 5’-GTCCGTGAAGAAACGATGATG-3’ and reverse primer 5’GAGCACTTACAAGCACGGG-3’, ViMs targeting exon 50 or 52: forward primer exon 49: 5’-AAACTGAAATAGCAGTTCAAGC-3’, reverse primer exon 54 5’-CCAAGAGGCATTGATATTCTC-3’). The PCR consisted of 36 cycles (40 sec at 95°C, 40 sec at 60°C and 90 sec at 72°C). Products were visualized on a 2% agarose gel.

## Results

The hDMD/*mdx* mouse model carries two tail-to-tail copies of the human *DMD* gene and remnants from the YAC in its integration site on mouse chromosome 5 (’t Hoen et al. 2008; Chey et al. 2024; Yavas et al. 2020). These complex genetics severely complicates the generation of exon deletions in the human *DMD* gene using genome editors (like CRISPR/Cas9, TALENs or ZNF-nucleases). Using editors on each side of the target exon, a large deletion between editor target sites could be induced, resulting in the loss of the 3’ end (downstream of the target exon) of both copies (Fig. 1). We optimized the PCR-based prescreen of edited samples based on this knowledge by combining a PCR assay spanning the predicted deletion, to confirm the presence of the intended allele and retention of the region 3’ from the target, with a PCR assay with primers outside and inside the deletion to score for the absence of the target exon. The first assay scores positive for correct deletion of the target exon, but false-positive if only one of the two copies of the target exon is deleted, while the second assay identifies these with a positive result. Combinedly, these two assays should efficiently identify the candidate clones or animals for further detailed testing (Fig. 1).

### Screening of the hDMDdel44/mdx model

We tested the revised prescreen by deleting exon 44 of the human *DMD* gene in embryonic stem (ES) cells that had previously been derived from the hDMD/*mdx* mouse model (Veltrop et al. 2013). CRISPR/Cas9 RNP complexes targeting introns 43 and 44 were used to delete human exon 44 from these cells, and clones were tested with the new prescreen assays. We found an unexpected high percentage of candidate clones (14%) passing both steps of the prescreen (Fig. 2). Not every clone with a del44-positive PCR fragment was the same length, suggesting different deletions in different clones. This is to be expected, as non-homologues end-joining repair will not always give the same outcome and can be biased to certain deletions by microhomology-dependent repair. Selected clones were further expanded, retested, and subjected to chromosome counts. A clone that passed the quality control criteria was used for blastocyst injection, chimera production and breeding for germline transmission to generate the hDMDdel44/*mdx* mouse model. Amplification and Sanger sequencing of the target region (Fig. 3A, Supplementary Table 2) showed that for the del44 model indeed only one deletion could be detected, suggesting that both copies of the YAC had obtained exactly the same deletion.

**Figure 3.**
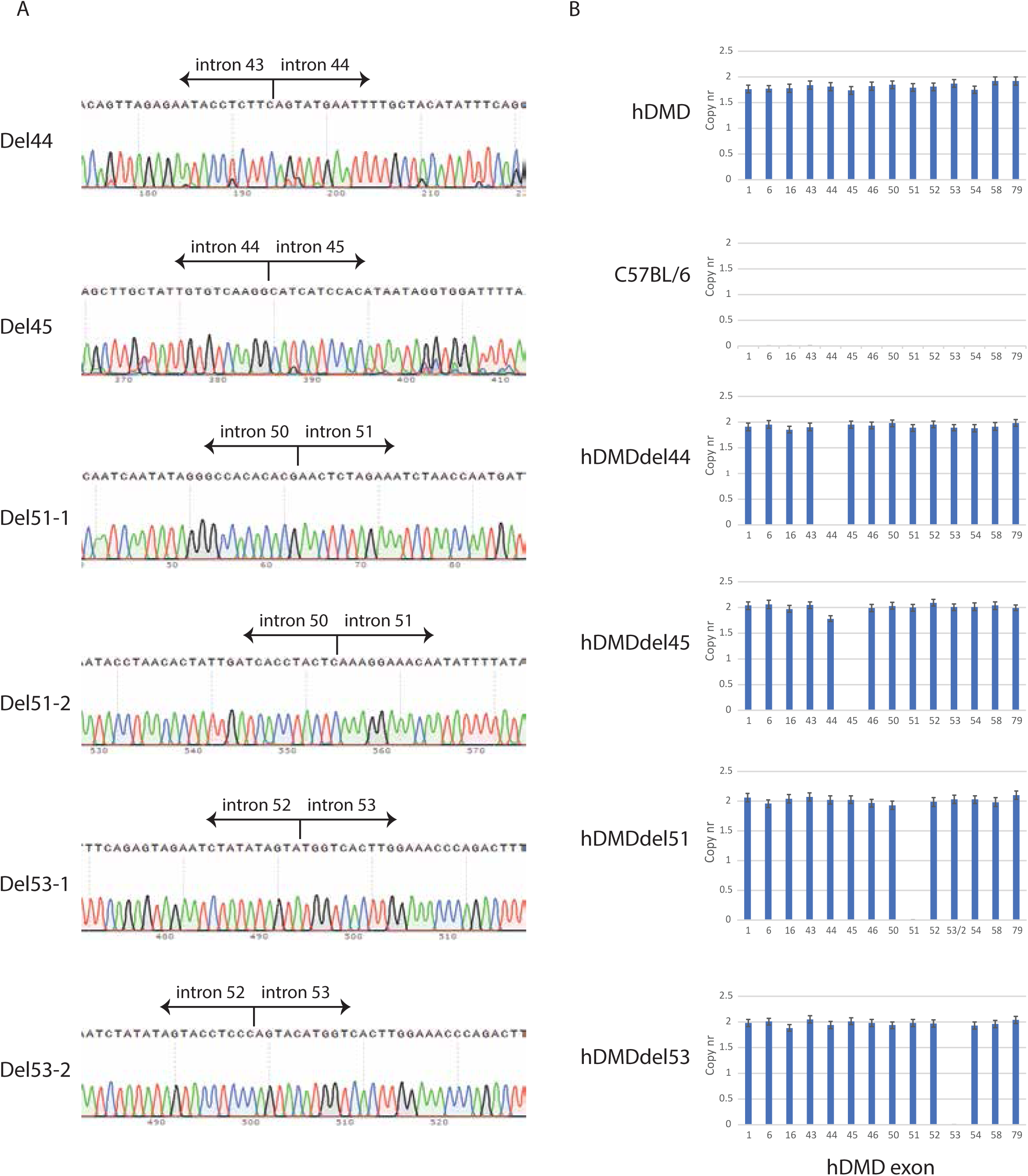
Additional quality control new hDMD models. **A.** Sanger sequencing of all mutant YAC alleles. del51-1 and del51-2 are the two genetic variants observed in the founder animals. del53-1 and del53-2 are the two different deletions in the two copies of the YAC. **B.** Copy counting of mutant and control exons using ddPCR.

### Screening of the hDMDdel45/mdx, hDMDdel51/mdx and hDMDdel53/mdx models

Encouraged by the specificity of our new prescreen workflow and the efficiency of correct targeting of human exon 44 in ES cells, we generated additional models with either a deletion of exon 45, 51 or 53 directly in mouse zygotes. Where possible, we used homozygous hDMD/*mdx* males and wildtype C57BL/6J females to generate zygotes where every embryo carried a single tail-to-tail hDMD insertion. Mutation-specific prescreen PCRs were used to identify candidate founder animals for the del45, del51 and del53 models.

Founder animals of the del45, del51 and del53 models were bred for germline transmission and mice hemizygous for two copies of the intended mutant YACs were readily obtained. Amplification and Sanger sequencing of the target region (Fig. 3A) showed that for the del45 model only one deletion could be detected, while for the del51 model, we initially only identified founder animals lacking one copy of the target exon, with the second copy still intact. We therefore selected a second set of single guide RNAs to completely delete exon 51. These single guide RNAs were located within the region already deleted in the first copy of the YAC, thus assuring that the already mutant YAC could not be edited further. Mice carrying the single del51 allele were used to generate new zygotes, which were targeted with the new single guide RNA set. In this second targeting round we were successful in identifying founder animals lacking both copies of exon 51 (Fig. 3A).

For the del53 model, we initially obtained only mixed Sanger sequencing traces, suggesting two different deletions in the two copies of the YAC. Indeed, TOPO cloning of the PCR product and sequencing of individual clones could separate the two deletions (Fig. 3A).

### Copy number analysis of the new hDMD deletion models

To test the integrity of the remaining parts of the YAC alleles in each model, we performed digital droplet PCR copy counts of selected exons spread over the totality of the human *DMD* gene, including the target exons and exons directly up- and downstream from the target exons. This analysis showed the presence of two copies for each exon (as expected from the double YAC integration in the original model) except for the target exon in each new model which was absent (Fig. 3B).

### Validation of a muscular dystrophy phenotype in the novel hDMD deletion models

The hDMD/*mdx* model and its deletion derivates are primarily of the C57BL/6J background. To reduce the amount of backcrosses required to generate a line with the *mdx* mutation, we crossed the models directly to the *mdx* line of the C57BL/6J background. The consequences of the exon deletion on dystrophin expression in the novel models was assessed by Western blot and immune fluorescence analysis on gastrocnemius muscles of 8-week-old males. As expected, the gastrocnemius of healthy hDMD/*mdx* mice expressed dystrophin of human origin at wildtype levels. Dystrophin expression in the hDMD deletion models was however either lacking (hDMDdel44/*mdx* and hDMDdel53/*mdx*), or trace dystrophin levels were observed (hDMDdel45/*mdx* and hDMDdel51/*mdx*) of human origin (Fig. 4A-B). The gastrocnemius of all novel deletion models was characterized by the classical histopathological hallmarks of the disease, comparable to those found in *mdx/*BL6 mice and consisting of muscle degeneration and regeneration, inflammation and fibrosis (Fig. 4B).

**Figure 4.**
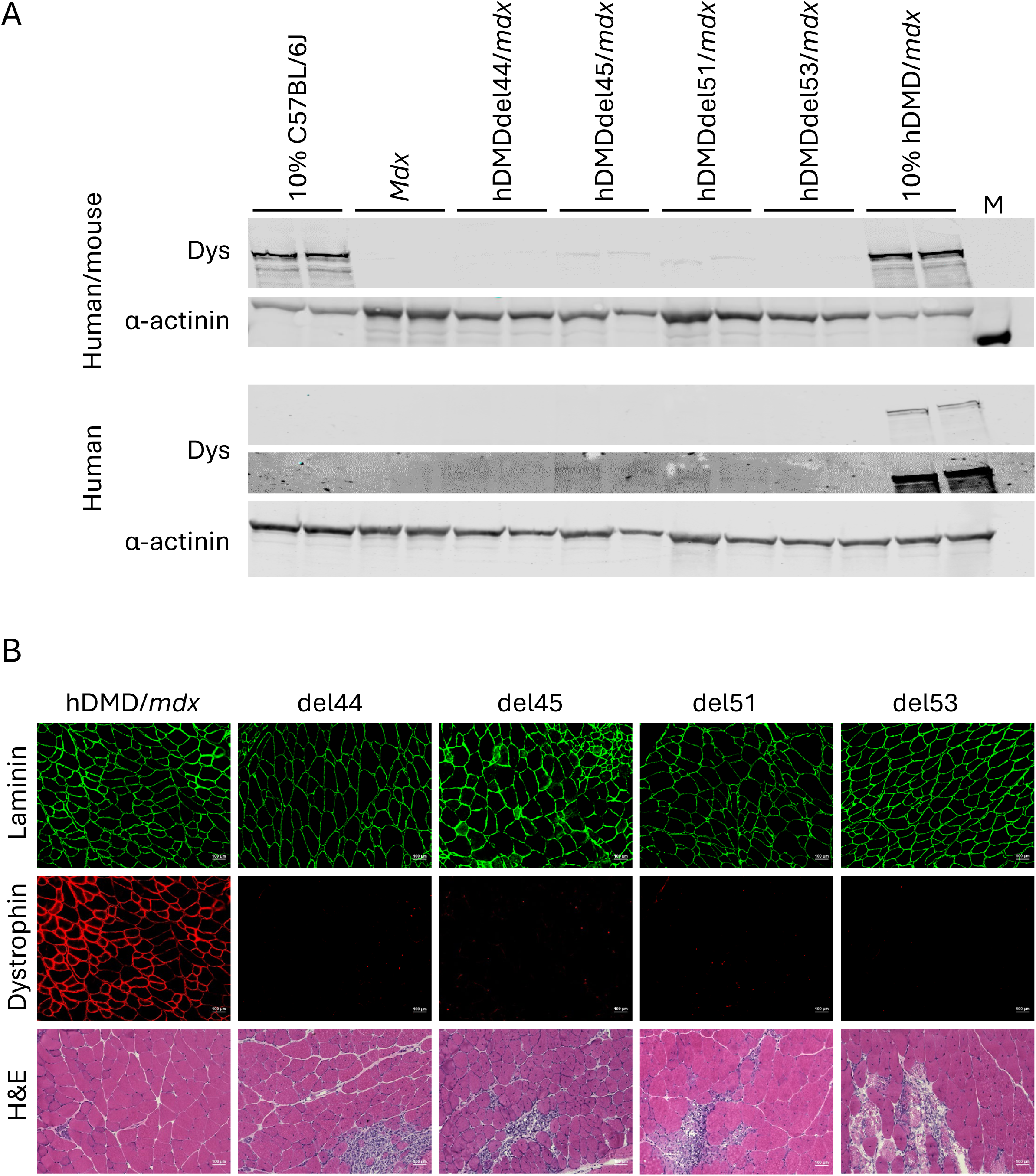
Characterization of dystrophin expression and histopathology. **A.** Dystrophin expression (of either human/mouse (top) or human specific origin (bottom, with a low and high exposure for dystrophin)) assessed by Western blot revealed that the hDMDdel44/*mdx* and hDMDdel53/*mdx* gastrocnemius completely lacked dystrophin expression, while traces of dystrophin were found in the hDMDdel45/*mdx* and hDMDdel51/*mdx* mice. Alpha-actinin served as loading control. M; marker. **B.** Laminin and dystrophin (top) and hematoxylin and eosin (H&E; bottom) staining in the gastrocnemius of 8-week-old mice. Barely detectable dystrophin positivity was observed in the hDMDdel51/*mdx* model. Histopathology consisted of de-, and regeneration, inflammation and fibrosis.

### Validation of the applicability of the hDMD deletion models for exon skipping

To validate that exon skipping can restore dystrophin expression in these models (thus ruling out the introduction of secondary mutations elsewhere in the human *DMD* gene during the editing process), we designed ASOs targeting exon 43 (for the hDMDdel44/*mdx* model), exon 44 (for the hDMDdel45/*mdx* model), exon 50 (for the hDMDdel51/*mdx* model) and exon 52 (for the hDMDdel53/*mdx* model). Either 100µg (exon 43 and 44) or 50µg (exon 50 and 52) ViMs was injected intramuscularly in the gastrocnemius and triceps of both sites on two consecutive days. Two weeks after the last injection muscles were isolated, and RNA and protein was isolated from muscle sections obtained from cryosections representative of the entire muscle. Skipping of the targeted exon was confirmed with RT-PCR in each of the exon deletion models (Fig. 5A). As expected, restoration of the disrupted open reading frame induced expression of truncated dystrophin proteins as assessed with Western blot analysis (Fig. 5B).

**Figure 5.**
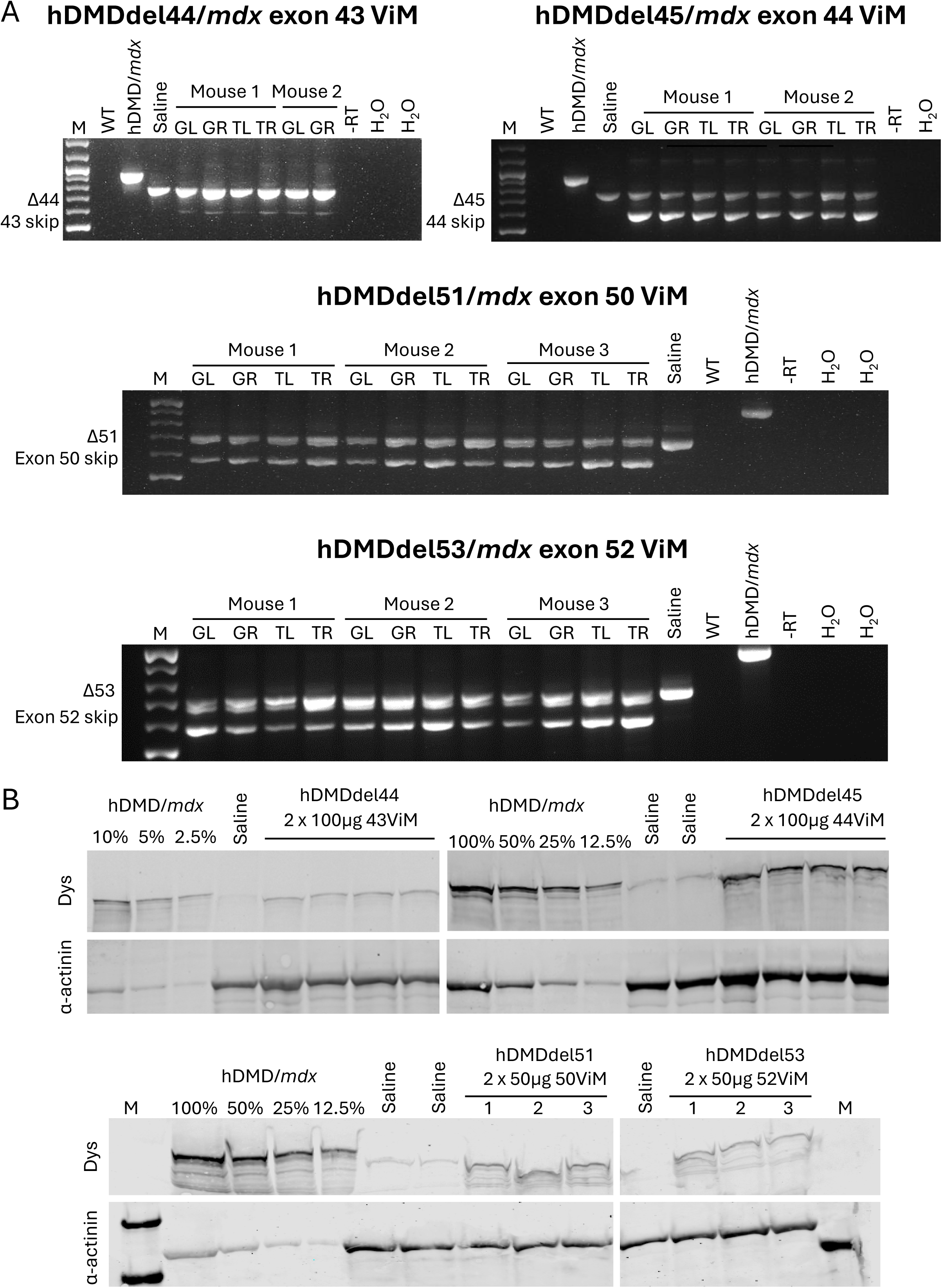
ASO-mediated exon skipping restores dystrophin expression in the deletion models. **A.** A single human specific PCR revealed skipping of exon 43, 44, 50 or 52 upon ViM injection of the right and left gastrocnemius and triceps muscles (respectively GR, GL, TR and TL) of either two or three hDMDdel44/*mdx*, hDMDdel45/*mdx*, hDMDdel51/*mdx* and hDMDdel53/*mdx* males. The upper (WT) band corresponds to the PCR product only lacking the deleted exon, while the lower (skipped) band additionally lacks the nucleotides of the skipped exon. M; 100bp ladder. -RT: no reverse transcriptase. WT; C57BL/6J sample. **B.** Western blot analyses confirmed that ASO-induced skipping of the respective exons restored the disrupted open reading frame and thereby exon skip expression in all the deletion models. A reference series of several dilutions of a wildtype sample was taken along as reference of the dystrophin levels restored.

## Discussion

Humanized mouse models are invaluable for preclinical testing of therapies that rely on the human DNA, RNA transcript or protein sequences of the causative mutated gene. We previously generated a humanized hDMDdel52/*mdx* model, which has since been widely used for preclinical studies (Van Deutekom et al. 2023; Engelbeen et al. 2023; Lim et al. 2022; Oppeneer, Qi, Henshaw, Larimore, Melton, et al. 2025; Oppeneer, Qi, Henshaw, Larimore, Puoliväli, et al. 2025). However, generating this model was cumbersome and while doing so we identified a double tail-to-tail integration of the human YAC in these mice (Yavas et al. 2020).

Here, we show that optimizing the prescreening strategy of genome edited ES clones and founder animals based on this knowledge leads to the efficient generation of new humanized patient-specific mutant DMD mouse models. We generated four new models for the recurring deletions of exon 44, 45, 51 and 53. These models either completely lack dystrophin, or express trace levels. This is in line with the trace dystrophin levels observed in DMD patients amenable for e.g. exon 44 skipping, and further increases the translational value of these models. The novel models all develop histopathology comparable to that found in the classic *mdx* model. Currently, work is ongoing to fully characterize the natural disease history of these newly derived models on motor function, histological and molecular levels throughout the lifespan in great detail. This will reveal whether there are actual differences between the models in disease development and severity and can further guide preclinical study design. Furthermore, we showed that ASO-mediated skipping of a flanking exon restores production of dystrophin, thus validating the models to be used to optimize mutation specific therapies.

Together with the hDMDdel52/*mdx* model, these new models cover the top four most frequently observed single exon deletions found in DMD patients (exon 45 (4%), 51 (3%), 44 (3%) and 52 (3%)), and could be utilized to tests single exon skipping ASOs that would combinedly be applicable for over 50% of mutations (Bladen et al. 2015). It is challenging to give an exact number as some mutations are amendable to skipping of either the exon before or after the deletion. For example, patients with a mutation of exon 44 are amenable for skipping of either exon 43 or 45.

With our optimized prescreening workflow it would be straightforward to create additional patient-relevant preclinical mouse models for human sequence-specific DMD therapies. The two copies of the YAC in our humanized model do not complicate the use of these models for testing ASO-mediated exon skipping preclinical testing. In contrast, the double integration makes the expression level in the mouse model more comparable to the human situation (’t Hoen et al. 2008). However, gene editing-based therapies could run into the same unpredictable outcome reduced efficiency of a therapy as we experienced when generating these mutant models, and preclinical outcomes could therefore be underestimated. Recently, Chey et al. sequenced the complete double integration site of the YAC transgenes in our hDMD/*mdx* model, and described guide RNAs that can be used to delete one copy of the YACs (Chey et al. 2024). This approach could be used to convert our mutant models into single-copy models as well, facilitating preclinical investigations on gene editing for DMD.

Taken together, we here present four novel dystrophic DMD mouse models that allow for preclinical testing of human sequence specific therapeutic approaches aimed to restore dystrophin expression. Utilization of these models could greatly enhance translatability of preclinical findings to the clinic.

## Acknowledgements

We would like to thank Jorrit Hos and Orhan Demirbacak for their experimental help. Regenxbio is kindly acknowledged for funding the vivo-morpholinos. This work was supported by a grant from AFM Telethon to Maaike van Putten (grant number 24745).

