## Supplementary Tables for "Four new Duchenne muscular dystrophy mouse models with clinically relevant exon deletions in the human *DMD* gene"

### Supplementary Table 1. Sequences guide RNAs

| Target exon | Upstream guide | Sequence | Downstream guide | Sequence |
|---|---|---|---|---|
| **44** | 110rev | AGCATAAAAAAGGCAACCGA | 1284forw | TTTCAATGATATCCAACCCA |
| **45** | hDmd_Ex45_upstr | TTGCTATTGTGTCAAGGAGT | hDmd_Ex45_downstr | CTATTATGTGGATGATGGGT |
| **51-1** | hDmd_Ex51_upstr | GTCAGTCAAAAGTCCGTGTG | hDmd_Ex51_downstr | TAGATTTCTAGAGTTCGTGG |
| **51-2** | hDmd_Ex51_2_upstr | ACAGCGCCTGACACTATAGT | hDmd_Ex51_2_downstr | CCAGGACTTTTATTTACCAA |
| **53** | hDmd_Ex53_upstr | TTAGGAGCTGTCCCCCAACA | hDmd_Ex53_downstr | TCATTGGAAGTCCAAGAGGG |

### Supplementary Table 2. Prescreening primers

| Exon | | Forward | Sequence | Reverse | Sequence | Size (bp) |
|---|---|---|---|---|---|---|
| 44 | del | hDMD int43-Fw1 | ACTCTTTGACTCCAGTGATGCAA | hDMD int44-Rev | TTCCAAGAAACCCAGCCTGA | 318 |
| | wt | hDMD-ex44/Fw1 | CCTGCAGGCGATTTGACAGA | hDMD int44-Rev | TTCCAAGAAACCCAGCCTGA | 262 |
| 45 | del | hd_Int44_2_Fw | TCACATGCAACGCTGATCTG | hd_Int45_1_Rev | GTGGGCAAAGACAACTGAC | 530 |
| | wt | hd_Int45.1_Fw | TGGCTAGTTAGTGGTTTTCTGC | hd_Int45.1_Rev | GTGGGCAAAGACAACTGAC | 700 |
| 51-1 | del | hd_Int50_2_Fw | GGTGTCCCTGAGAAGCTTGA | hd_Int51_4_Rev | AGTAGAGATGGGGTTTGGCC | 880 |
| | wt | hd_Int50.1_fw | AGCCTGGTGTTCTGCAGTTA | hd_int50.1_Rev | TGAGTGGTGAGGGGAAGGAA | 730 |
| 51-2 | del | hd_Int50_2_Fw | GGTGTCCCTGAGAAGCTTGA | hd_Int51_5_Rev | CCTGATGACAAACTGCAAGG | 800 |
| 51-1 and 51_2 | del | hd_Int50.3Fw | CATTCCTTCCCCTCACCACT | hd_Int51.3_Rev | TGGCTTCCCTAAATGCTGGA | 300 copy 1<br>730 copy 2 |
| | wt | hd_Int50_4_Fw | GGCATTGTCATACGTGTATTGCT | hd_Int50.4_Rev | CCCTAGGGAAATCAAAGCCAATG | 530 |
| 53 | del | hd_Int52_1_Fw | AGAGGGAGTGCAGTGATCAG | hd_Int53_1_Rev | TGTCCTCATAGCAGCTCAGG | 700 |
| | wt | hd_Int52.3_Fw | TGGTGCATTATCAAGCCAGC | hd_ex53.1 Rev | AGACCTGCTCAGCTTCTTCC | 757 |

### Supplementary Table 3. ddPCR primers and probes

| **Exon** | **Probe** | |
|---|---|---|
| **1** | hDMD-ex1/P1 | TGTTGGGATCACTCACTTTCCCCC |
| **6** | hd_Ex6_Probe | TGGCTGGATTGCAACAAACCAACA |
| **16** | hd_Ex16_Probe | AGCAATCCATGGGCAAACTGTATTCA |
| **43** | hDMD-ex43/P1 | TGCAACGCCTGTGGAAAGGGT |
| **44** | hDMD-ex44/P1 | TCTGTTGAGAAATGGCGGCGT |
| **45** | hDMD-ex45/P1 | TCAGCAATCCTCAAAAACAGATGCCA |
| **46** | hd_Ex46_Probe | CCACTTGAACCTGGAAAAGAGCAGC |
| **50** | hd_Ex50_Probe | ACTGGATCCCATTCTCTTTGGCTCT |
| **51** | hd_Ex51_Probe | CAACGAGATGATCATCAAGCAGAAGGT |
| **52** | hd_Ex52_Probe | ACAGAGGCGTCCCCAGTTGGA |
| **53** | HD_Ex53_P2 | TCAGAACCGGAGGCAACAGTTGAA |
| **54** | hd_Ex54_Probe | CTGCAGATGATACCAGAAAAGTCCACA |
| **58** | hd_Ex58_Probe | ACTCTACCAGGAGCCCAGAGGT |
| **79** | hDMD-ex79/P1 | GGATTTTTCCCGGAGCCGGAA |
| **Exon** | **Forward** | |
| **1** | hDMD-ex1/F1 | GGCCTCTACAGAATCCTGGC |
| **6** | hDMD-ex6/F1 | GGGTCCGACAATCAACTCG |
| **16** | hd_Ex16_Fw | TGGAATGCAACCCAGGCTTA |
| **43** | hDMD-ex43/F1 | AGCAAGAAGACAGCAGCATTG |
| **44** | hDMD-ex44/F1 | CCTGCAGGCGATTTGACAGA |
| **45** | hDMD-ex45/F1 | AGAACATTGAATGCAACTGGGG |
| **46** | hd_Ex46_Fw | TGGTTGGAGGAAGCAGATAACA |
| **50** | hd_Ex50_Fw | ACCTAGCTCCTGGACTGACC |
| **51** | hd_Ex51_Fw | GTGATGGTGGGTGACCTTGA |
| **52** | hd_Ex52_Fw | ACACAACGCTGAAGAACCCT |
| **53** | HD_ex53/F2 | GGGATGAAGTACAAGAACACCT |
| **54** | hd_Ex54_Fw | TTGGCCCTGAAACTTCTCCG |
| **58** | hd_Ex58_Fw | CAGAGCAGCCTTTGGAAGGA |
| **79** | hDMD-ex79/F1 | GGCTTACCTGCTTGGTCTAGAAT |
| **Exon** | **Reverse** | |
| **1** | hDMD-ex1/R1 | GCCTCCCAGATCTGAGTCCT |
| **6** | hDMD-ex6/R1 | CTATGGATGAGAGCATTCAAAGCC |
| **16** | hd_Ex16_Rev | CATGCTTCCGTCTTCTGGGT |
| **43** | hDMD-ex43/R1 | AGCTGGGAGAGAGCTTCCTG |
| **44** | hDMD-ex44/R1 | TCAGCTTCTGTTAGCCACTGA |
| **45** | hDMD-ex45/R1 | CGCAGATTCAGGCTTCCCAA |
| **46** | hd_Ex46_Rev | CTTCTTTATGCAAGCAGGCCC |
| **50** | hd_Ex50_Rev | CTCTCACCCAGTCATCACTTCA |
| **51** | hd_Ex51_Rev | GTTGCCTAAGAACTGGTGGGA |
| **52** | hd_Ex52_Rev | TTGGGCAGCGGTAATGAGTT |
| **53** | HD_ex53/R2 | CCTGCTCAGCTTCTTCCTTAG |
| **54** | hd_Ex54_Rev | CATCCAGTTTCACCACCCCA |
| **58** | hd_Ex58_Rev | AAGAATGGATGGGCTGCTCC |
| **79** | hDMD-ex79/R1 | AGTGTGGTGTAGTTTCCTCCTGG |

### Supplementary Table 4. ViM sequences

| **Mouse strain** | **Target exon** | **ViM sequence 5'-3'** |
|---|---|---|
| hDMDdel44/*mdx* | 43 | TTGTTAACTTTTTCCCATTGGAAAT |
| hDMDdel45/*mdx* | 44 | CTTAACCCTTGTACGATTTATGTTT |
| hDMDdel51/*mdx* | 50 | CCTTCCACTCAGAGCTCAGATCTTC |
| hDMDdel53/*mdx* | 52 | GCGGTAATGAGTTCTTCCAACTGGG |
